# Richness and density jointly determine context dependence in bacterial interactions

**DOI:** 10.1101/2023.05.02.539006

**Authors:** Keven D. Dooley, Joy Bergelson

**Author notes:** **Email addresses of corresponding authors** Correspondence. Further information and data requests should be directed to the lead contact, Keven D. Dooley.

## Abstract

Pairwise interactions are often used to predict features of complex microbial communities due to the challenge of measuring multi-species interactions in high dimensional contexts. This assumes that interactions are unaffected by community context. Here, we used synthetic bacterial communities to investigate that assumption by observing how interactions varied across contexts. Interactions were most often weakly negative and showed clear phylogenetic signal. Community richness and total density emerged as strong predictors of interaction strength and contributed to an attenuation of interactions as richness increased. Population level and per-capita measures of interactions both displayed such attenuation, suggesting factors beyond systematic changes in population size were involved; namely, changes to the interactions themselves. Nevertheless, pairwise interactions retained some predictive value across contexts, provided those contexts were not substantially diverged in richness. These results suggest that understanding the emergent properties of microbial interactions can improve our ability to predict features of microbial communities.

## Introduction

Microbes are the engines of many biochemical processes that support life on Earth^1^. Importantly, however, microbes rarely perform these complex functions in isolation, instead acting within communities. Many efforts are thus underway to design microbial communities that perform desired functions, enabling us to co-opt these powers of chemical transformation and develop applications relevant to human health, agriculture, and industry^2,3^. However, the intricate relationships underlying such complex functions provide a challenge that must be overcome, as interactions among members constrain the extent to which the abundance and distribution of a focal microbe can be manipulated. Overcoming this challenge will require understanding the forces that determine the structure and function of microbial communities.

Interactions between community members have long been known to affect community composition^4,5,6^ and therefore the emergent functions performed by a community^7,8^. Leveraging an understanding of interspecific interactions is a promising and actively researched approach for designing the structure and function of microbial communities^9,10^. However, for such an approach to be effective, observations of interactions made in one community context must inform the extent of that interaction in another context.

Interactions are often modelled as a network of static pairwise per-capita or proportional effects between members of a community^11,12,13^. By assuming that it is appropriate to distill an interaction into a simple static relationship, we can reduce the complexity of interaction networks^14^ and apply knowledge of interactions gleaned from other contexts to make predictions about unobserved communities^15^. However, a variety of known effects call this simplification into question. Interactions can be subject to higher order effects (“higher order interactions” or “HOIs”) where a pairwise interaction is altered by the presence of one or more other community members^16,17,18^. Habitat modification can also affect microbial interactions^19^, an example being environmental pH modification, which has been observed as a relevant factor in microbial community assembly^20,21,22^. Due to effects such as these, knowledge of pairwise interaction strength or coexistence can have limited predictive power in complex communities^23,24^. Thus, advancing our understanding of what contributes to the variation of interactions between contexts stands to facilitate the rational design of microbial communities.

One potential solution to these complexities is to identify patterns in how pairwise interactions vary across contexts and uncover the underlying drivers of this variation. Such an understanding stands to improve our predictions of how microbial interactions will change between community contexts. Encouragingly, recent work has demonstrated that stronger negative interactions are found at high nutrient concentrations^22^, confirming the possibility of identifying broadly general patterns. By expanding our understanding of such patterns, we hope to improve the predictive power of pairwise interactions. Here, we use synthetic bacterial communities to observe how interactions vary across community contexts and identify patterns underlying that variation.

## Results

### Assembly of synthetic communities

We assembled a set of synthetic communities from a pool of 56 bacterial strains isolated from the leaves of wild and field-grown *Arabidopsis thaliana* by randomly dividing isolates into seven pools of eight members. We then created 127 unique communities representing all possible combinations of those pools (i.e., seven single pool communities, twenty-one two pool communities, etc.) (figure 1a). These communities were inoculated into a custom growth-medium derived from *A. thaliana* leaves (*<u>A</u>rabidopsis* leaf <u>m</u>edium, ALM) (STAR methods) at a consistent total community titer, with each member accounting for an equal proportion of the population given the initial richness (number of community members). To allow the communities to reach a steady state reflective of its long-term composition, we passaged each community for 6 days by performing a 1:100 dilution into fresh medium every 24 hours (figure 1b). This period was sufficiently long to allow the community composition to stabilize (supplementary figure 1). We characterized the compositions of these final communities by mapping Illumina short reads against a nearly complete and high-quality genome of each isolate (STAR methods).

**Figure 1.**
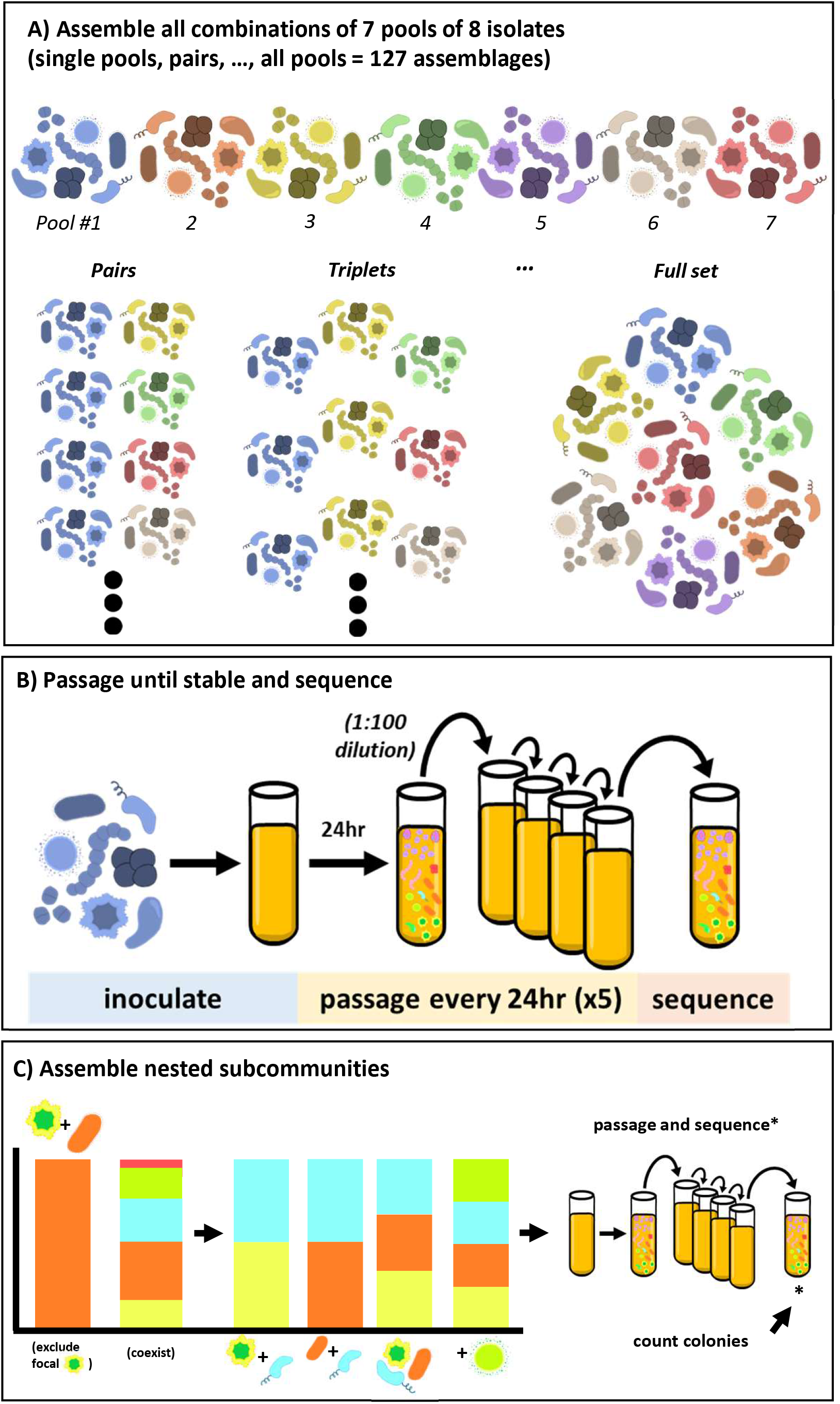
Experimental outline: A) A set of 56 isolates representing 21 genera were randomly pooled into 7 pools. All combinations of those pools were assembled at equal titers, with respective densities scaled to the total number of isolates initially present. B) These combinations were inoculated in triplicate into a custom medium derived from *Arabidopsis* leaves (ALM) and passaged daily into fresh medium at a 1:100 dilution for 5 days. To characterize the community compositions, the day-6 samples were sequenced, and short-reads were mapped to reference genomes. C) Ten communities displaying context-dependent coexistence were decomposed into nested subcommunities containing the focal isolate and/or putative excluder isolate. These communities were assembled, passaged, and sequenced as described for the previous communities. To provide the absolute abundance information necessary to measure interactions, the final timepoint (day-6) was quantified by counting colonies on 1X TSA plates.

### Measurement of interactions

We screened this set of 127 communities for putative interactions by finding pairs of communities where a focal isolate was observed to coexist alongside one or more specific isolates in one community context but was excluded in another context (figure 1c, supplementary figure 2). We posited that such context-dependent coexistence was related to interactions between the focal isolate and/or its context-dependent excluder with additional members of the community. Thus, from all paired communities in which we observed context-dependent coexistence, we selected a set of ten pairs that maximized compositional diversity in which to investigate potential interactions.

To do so, we decomposed these communities into sets of nested subcommunities varying by a single isolate (figure 1c), always including a focal isolate and/or its excluder. In total, we assembled 245 such communities and passaged them for 6 days, as previously described. A subset of communities was passaged for 12 days, with samples from days 6 and 12 sequenced to confirm that community composition was stable by day 6 (supplementary figure 3). With these sets of nested communities, we were able to measure the effect of one isolate on the abundance of another (i.e., an interaction) across multiple community contexts (figure 2a).

**Figure 2.**
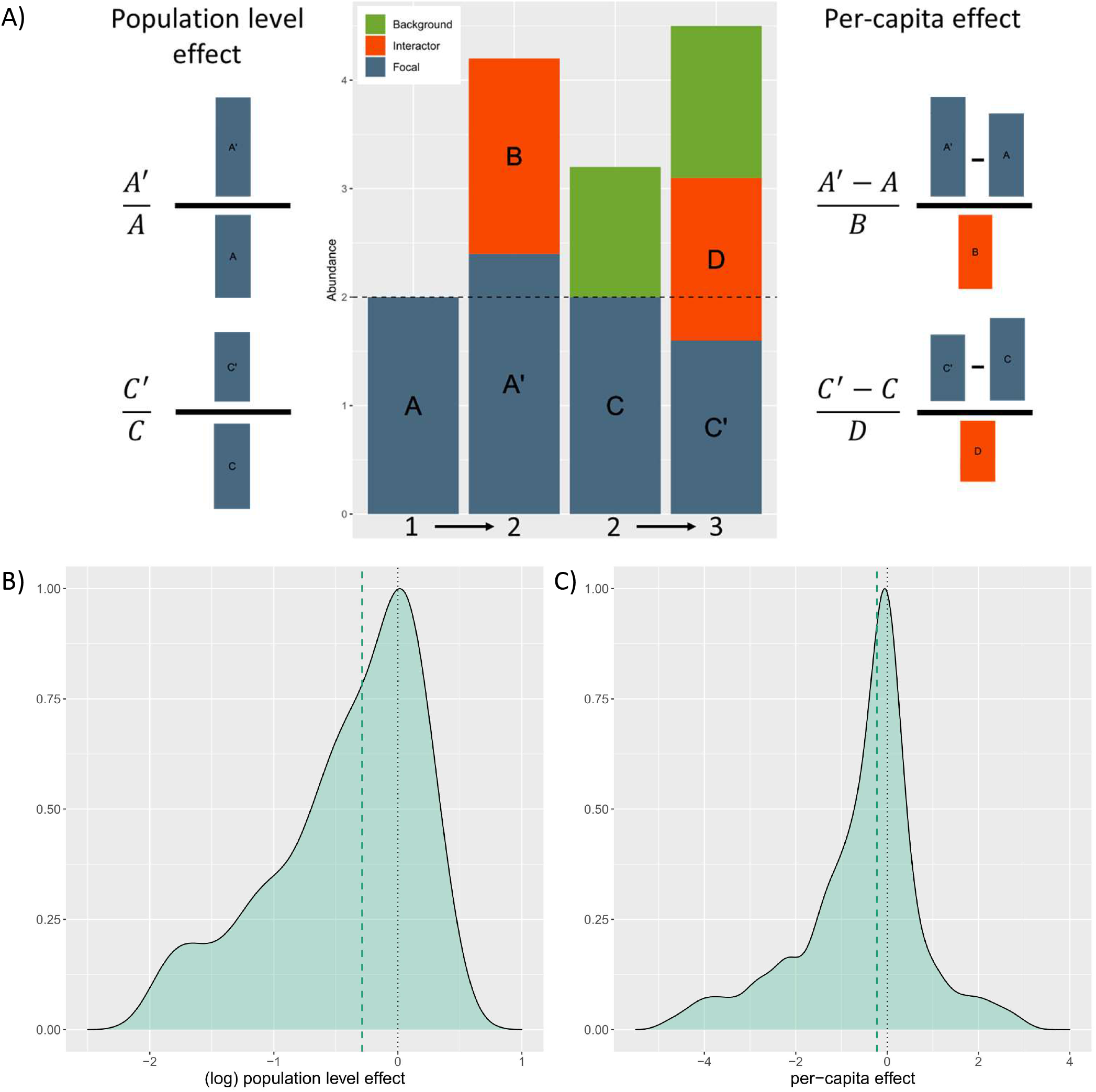
Distributions of observed interactions: A) Interactions between a “focal” isolate and “interactor” isolate were calculated as two measures, 1) a population effect, calculated as the ratio of the focal isolate’s density with and without the interactor present, and 2) a per-capita effect, calculated as the change in density of the focal isolate between contexts with and without the interactor, scaled by the abundance of the interactor. Interactions were always calculated between communities varying by a single isolate – the interactor. However, additional isolates (“background” isolates) could also be present in the compared communities. The “richness context” of an interaction refers to the richness of the pairs of community contexts from which an interaction is observed (e.g., 1=>2 for the first example interaction, 2=>3 for the second example interaction including a “background” isolate). C) the distribution of all observed interactions, as population level effects, (natural-log transformed to symmetrize ratios), C) the distribution of all observed interactions, as per-capita effects.

Importantly, interactions are measured as changes in absolute abundance, but characterizations of community composition obtained through sequencing are limited to relative abundance information. Thus, we measured absolute abundances by counting colonies from serial dilutions of each day-6 sample (STAR methods), which we then used to translate the relative abundances obtained from sequencing into absolute abundances.

With measures of absolute abundance in hand, we measured interactions by comparing abundances between pairs of communities that varied by a single member (the “interactor”). For example, the interaction between a “focal” isolate A and an “interactor” isolate B was observed by comparing the abundance of A in a community lacking B to the abundance of A in a community where B was present. Here, we measure interactions using two metrics (figure 2a, STAR methods), and refer to the signed effect of an interaction as its “strength” and the absolute value as the “absolute strength”. Our first metric measures an interaction as the ratio of the focal isolate’s abundance in the context with the interactor to its abundance in the context without the interactor. This is a commonly used metric^25,26,27,28,29^, which represents an interaction as a population level effect on the focal isolate. Our second metric measures an interaction as the per-capita effect of an interactor on the abundance of a focal isolate^14^. The population level effect of an interaction is a function of the per-capita effect of an interactor scaled by the density of that interactor in a given community context. This is relevant, as below we will show that a general relationship between richness and density existed in our communities and contributed to the observed effect of interactions.

### Negative interactions were more common and stronger than positive interactions

We observed a total of 388 pairwise interactions across all community contexts (figure 2). Negative interactions were more common, representing 67% of population level interactions. We observed median values of -0.29 for the population level and -0.22 for the per-capita effects, respectively (figure 2). Negative interactions were statistically greater in absolute strength than positive interactions for both measures (one-tailed Wilcoxon rank sum test: p-values <4e^-8^ and 0.002, respectively). We also observed phylogenetic effects for both population level and per capita level measures of interactions. Namely, isolates belonging to the same genus tended to have a more negative interaction than those belonging to distinct genera (one-tailed Wilcoxon rank sum test: p-value 0.004 and 0.17, population and per-capita, respectively).

### Individual interactions attenuated as richness increased

Our method of measuring interactions was to compare contexts that differ in composition by a single “interactor” isolate. To observe how interactions changed between contexts, we compared the strength of pairwise interactions measured in community contexts with or without a single “background” isolate. In this way, we compared interactions between two pairs of communities that differed in “richness context” by one (figure 2a). As an example, we might compare an interaction measured when only the focal and interactor are present (richness context 1->2) with that interaction measured when a single “background” isolate was also present in both contexts (richness context 2->3). Use of our complete dataset enabled us to analyze 275 instances of such paired contexts. First focusing on the population level effects, we observed that interactions generally attenuated in strength when measured in a community with one additional background member (median difference -0.08, paired Wilcoxon signed rank test, p-value 0.003). When grouping interactions by their initial direction, the median positive and negative interaction became less positive and negative, respectively (figure 3a). Absolute strength was significantly weaker for initially negative interactions but significantly stronger for initially positive interactions (paired Wilcoxon rank sum test: p-values <4e^-7^ and 0.006, respectively). This result for the initially positive interactions is manifest as a shift from a weakly positive median interaction to a moderately negative median interaction.

**Figure 3.**
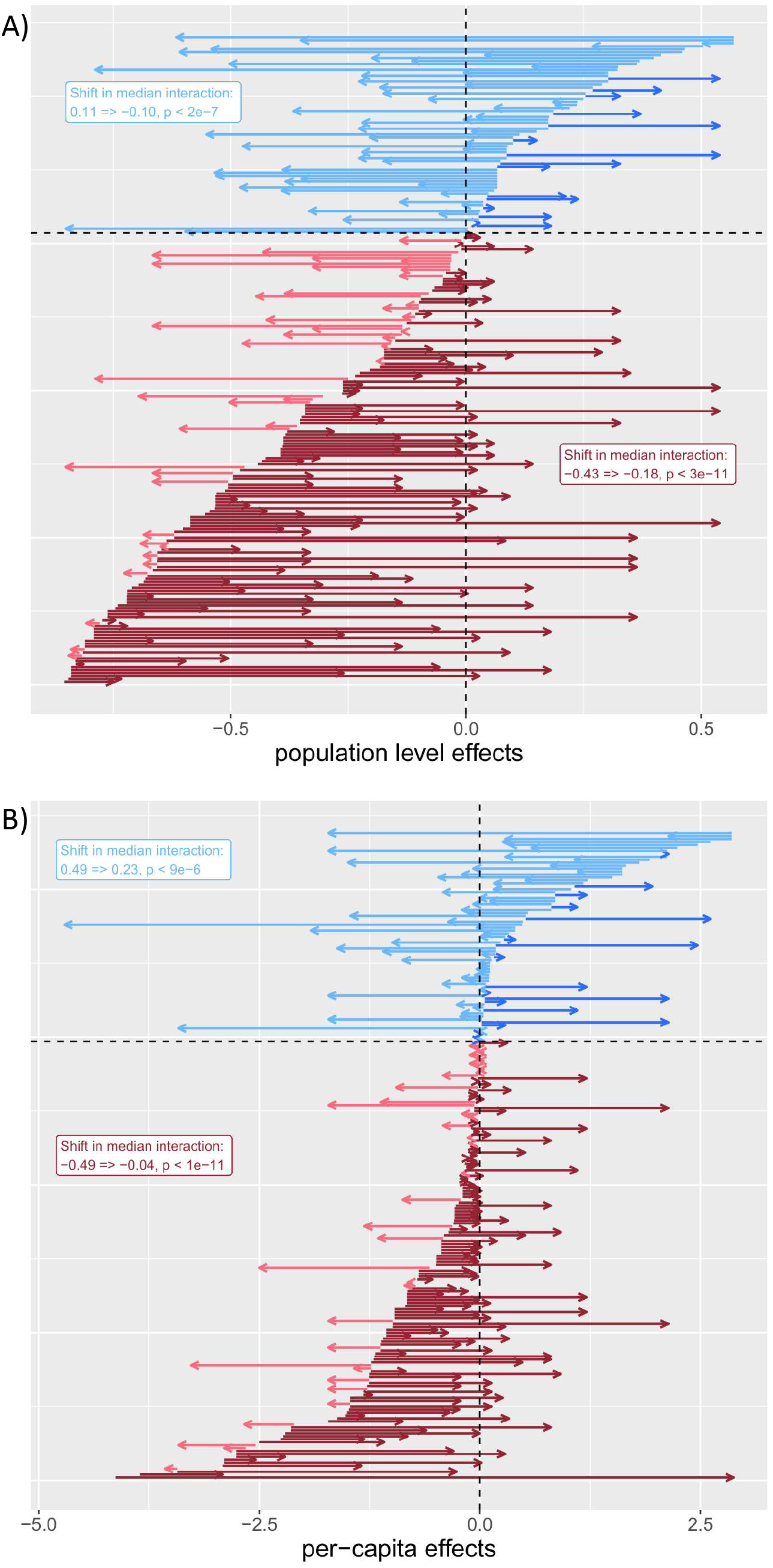
Interactions attenuated as richness increased: The shift in interactions between richness contexts varying by a richness of one for A) population level, and B) per-capita effects. All comparisons of an interaction that could be compared between richness contexts varying in richness by a single isolate. Each arrow on the plots represents an interaction observed in two separate richness contexts (e.g., 1=>2 and 2=>3 community members), with the tail of the arrow representing the value of the interaction in the lower richness context and the head of the arrow representing the value observed in the higher richness context. Arrows are colored by the initial direction of the interaction (positive or negative) and the shift in direction of the interaction (positive or negative). Labels display the shift in median interactions for initially positive and negative interactions, respectively, with p-values summarizing the outcome of Wilcoxon signed-rank tests to determine if the shift in interaction value represented a significant change.

As previously stated, the population level effect of an interaction is a function of the per-capita effect of an interactor and the density of that interactor in each community context. Thus, the observed decrease in population level effects suggests a decrease in the strength of per-capita effects and/or a systematic decrease in interactor density. Indeed, when we consider the shift in per-capita interactions, we observe again that interaction strength attenuated as richness context increased (Wilcoxon signed rank test, p-value 0.003). As with the population level effects, per-capita effects grouped by initial direction showed consistent shifts (figure 3b). Absolute strength became significantly weaker for initially negative interactions; however, unlike at the population level, it remained statistically unchanged for positive interactions (one-tailed Wilcoxon rank sum test: p-values 0.002 and 0.64, respectively). These results suggest that part of the decrease in the population level effects can be explained by a decrease in the per-capita effects. Next, we evaluate an alternative explanation by investigating the relationships between richness and density in our communities.

### Relationships between richness and density help explain trends in population effects across richness

We observed that, as richness increased, average total density of communities gradually increased to a modest extent (figure 4a), while the average density of each member decreased before reaching an asymptote in communities with four members (figure 4b). This relationship between richness and the density of community members was often observed at the individual isolate level as well (supplementary figure 4). This general decrease in density with increasing richness helps explain the observed attenuation of interactions when measured as a population effect. Namely, because individual densities decreased with an increase in community richness, the population level effect of the interactor should decrease. Indeed, interactor density explained a significant portion of variance in population level effects (linear regression, adjusted R^2^: 0.09, p-value <5e^-10^). Additionally, the relationship between the densities of community members and total community density meant that as richness increased, the absolute change in total density associated with an interaction decreased (figure 4c, Pearson’s r: -0.23, p-value 0.002). In other words, adding a given interactor to a community resulted, on average, in a smaller change to total community density in higher richness contexts.

**Figure 4.**
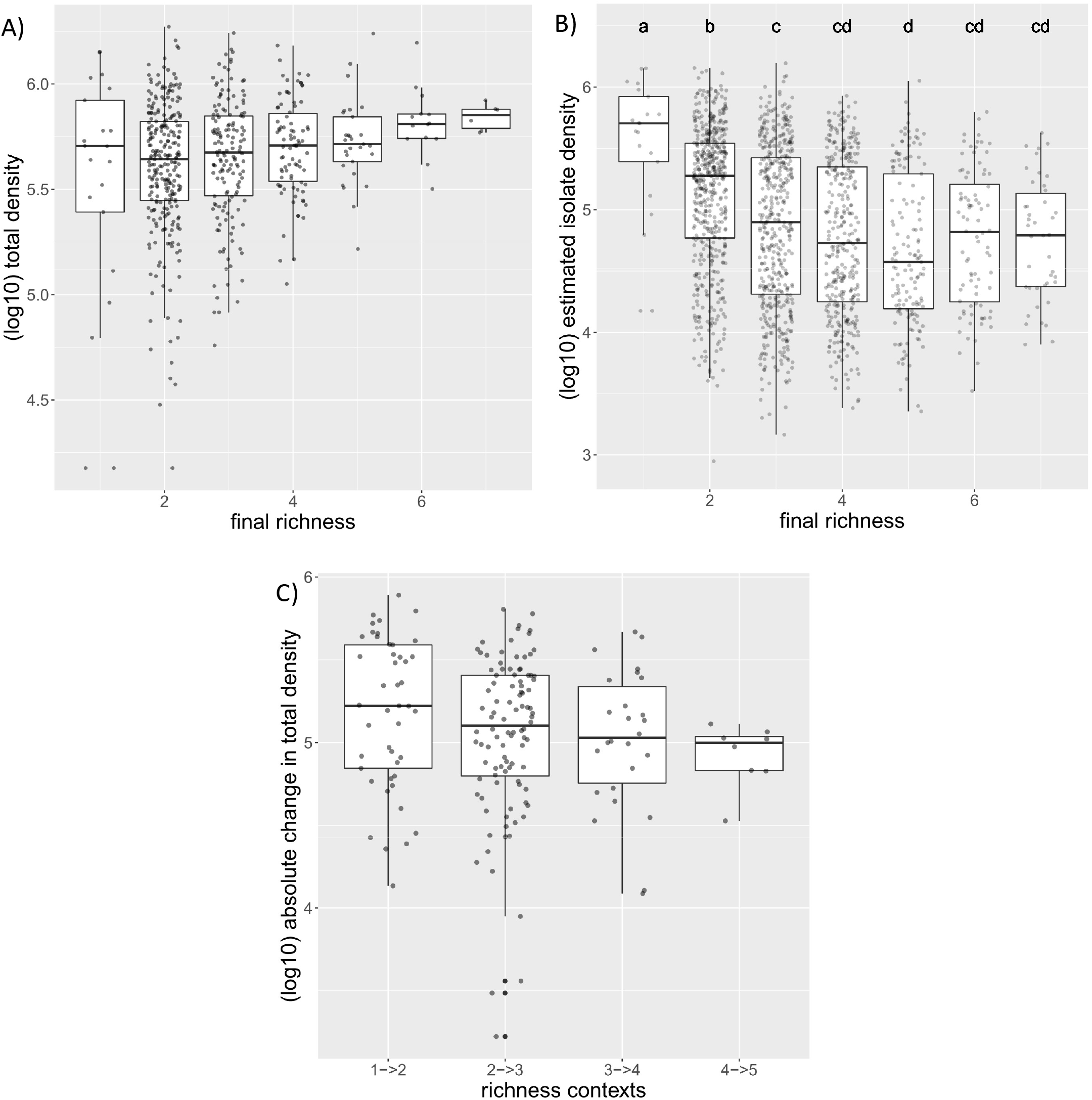
Relationships between richness and density: A) the relationship between final community richness and total community density, B) the relationship between richness and individual isolate density (significantly distinct groups were determined through post hoc pairwise t-tests using the Holm method for multiple testing correction), C) the absolute change in total community density associated with an interaction, grouped by the richness context in which interactions were observed (interactions from richness contexts with fewer than 3 observations were removed). For all plots, density is plotted on a log scale.

### Interactions are predictive between contexts

We next asked, what is the remaining predictive capacity of an observed interaction across community contexts? We attempted to answer this question by modelling the change in abundance of a focal isolate between community contexts (i.e., the effect of an interaction), informed by richness, total density, and interactions observed in different contexts.

First, while modelling the change in abundance associated with all 388 observed interactions, both richness and the change in total density emerged as highly explanatory variables associated with the change in focal isolate abundance (table 1). The change in total density was more explanatory with an adjusted R^2^ value of 0.33 compared to 0.08 for richness. A joint model including both variables and their interaction could explain 57% of variance in focal isolate change in abundance.

**Table 1.** Summary of linear regressions modelling the effect of an interaction by emergent community properties: “Focal change” indicates the change in density of the focal isolate in the predicted context. “Total change” indicates the change in total density between the community contexts of the interaction. “n-context” indicates the richness contexts over which the interaction was observed (as a factor). A “*” in the model indicates an interaction term in addition to the separate effects.

| Model | df | adjusted R<sup>2</sup> | p-value |
| --- | --- | --- | --- |
| focal change ~ total change | 1 | 0.334 | $< 2.2e^{-16}$ |
| focal change ~ n-context | 5 | 0.083 | $2.2e^{-7}$ |
| focal change ~ total change * n-context | 11 | 0.565 | $< 2.2e^{-16}$ |

Next, we used the set of 275 paired interaction contexts differing by a richness of one to interrogate the predictive power of interactions across contexts (e.g., compare interaction effects between richness contexts 1=>2 and 2=>3). In this dataset, interactions in one context were able to describe ∼16% of the variance in the change in density of a focal isolate (i.e., interaction effect) in the other context (table 2). A model using the change in total density of the context for which the interaction effect was being predicted (akin to the models above) explained ∼27% of variance. A joint model of these two variables (change in total abundance of the predicted context and interaction effect in another context) explained ∼42%, suggesting that the two variables are largely independent. Expanding the dataset to consider comparisons between any observations of a given interaction (e.g., compare richness contexts 1=>2 and 4=>5) reduced the explanatory power of interactions between contexts to ∼10% of variance, suggesting the predictive power of interactions decays as communities diverge in richness and composition (supplementary table 2). Ultimately, these results demonstrate the persistent but limited predictive power of interactions across contexts and highlight the relevance of community level properties in understanding the assembly of microbial communities.

**Table 2:** Summary of linear regressions modelling the predictive power of interactions between contexts differing in richness context by a single community member: “Focal change” indicates the change in density of the focal isolate in the predicted context. “Total change” indicates the change in total density between the community contexts of the interaction. “n-contexts” indicates the richness contexts from the pair of interactions (e.g., 1=>2 & 2=>3). “Interaction effect” indicates the change in density of the focal isolate in the interaction context which was not being predicted. We considered predictive power of interactions from the bottom-up, i.e., “interaction effects” came from the lower richness context (continuing the example above, 1=>2), while “total change” came from the predicted higher richness context (2=>3), as in the models described in Table 1. “+” models separate variables with no interaction, A “+” in the model indicates the separate effects with no interaction term.

| Model | df | adjusted R<sup>2</sup> | p-value |
| --- | --- | --- | --- |
| focal change ~ total change | 1 | 0.273 | $< 2.2e^{-16}$ |
| focal change ~ n-contexts | 4 | 0.023 | 0.035 |
| focal change ~ interaction effect | 1 | 0.159 | $< 2e^{-12}$ |
| focal change ~ total change + interaction effect | 2 | 0.417 | $< 2.2e^{-16}$ |

## Discussion

Here, we used synthetic bacterial communities to observe a large set of interactions across community contexts, ranging from the simplest possible community of two coexisting isolates to complex communities with up to seven isolates. These interactions were, on average, weakly negative and displayed a phylogenetic effect, in alignment with other studies of microbial interactions^27,28,30^. However, positive interactions were not uncommon, an observation that has growing recent empirical support^25,27^. When comparing interactions across contexts of increasing richness, we observed a general attenuation of interactions, though this arose predominately due to a consistent shift in negative interactions (figure 3). We observed that much of this change can be explained by relationships between individual density and richness/total density (table 1, figure 4). Namely, as richness increased, the modest increase in total density resulted in a decrease in individual isolate density (figure 4b). This relationship can help explain the observed attenuation of population level effects, as decreased density of interactor isolates in higher richness contexts should lead to smaller effects and did, in fact, explain ∼9% of variance in population level effects. However, the per-capita effects also showed some decrease in strength with an increase in richness, at least for negative interactions, suggesting additional processes were present that imparted a systematic change on the interactions.

Why would per capita effects be attenuated at high richness, and why predominately among initially negative interactions? Previous observation of the attenuation of pairwise interactions in the zebrafish gut was attributed to the effect of higher order interactions^31^, though that study was unable to identify the mechanisms of such effects. We have a similarly limited mechanistic understanding of observed interactions and what underpins their variation between community contexts. The importance of HOIs in microbial community assembly remains an actively debated subject, with theoretical and empirical evidence to support both sides^7,18,32,33,34,35,36^. However, HOIs are challenging to appropriately identify^16,37,38^, and our lack of fully characterized interaction networks precludes us from determining their relevance here.

Another possible explanation for the attenuation of per-capita effects is non-additivity in interactions. In other words, overlap in the mechanisms underpinning how multiple interactors affect a given focal isolate could result in a reduced per-capita effect when multiple interactors are present. Such non-additivity has been recently reported^25^. This effect would be likely if metabolic interaction (such as competition over labile carbon sources) predominately underlies interactions and community assembly, as has been shown in synthetic communities that were organized into functional guilds by preferred metabolic strategy^39,40,41^. In the context of how we measure and compare interactions here, such mechanistic overlap would be hypothesized to reduce the impact of a novel interactor due to a function already being performed by a “background” isolate in the community. Such mechanistic redundancy would be probabilistically more likely as richness increases.

Despite the limitations of our data, some insights can be inferred by asking what gives rise to the relationships we observed between richness and individual or total density. The apparent modest increase in total density in higher richness communities might have emerged for two reasons: 1) larger initial pools of isolates entailed greater metabolic diversity, thus allowing the community to occupy more of the available niche space, or 2) larger initial pools may simply have had a greater chance of including one or more isolates with high fitness in the environment (a “sampling effect”)^42^. Both possibilities would result in higher levels of community metabolic activity at higher levels of richness, which has been observed to have a positive effect on those community members with relatively low fitness as a result of cross-feeding or general metabolic leakiness^27,43,44^. In this way, positive effects absent in simpler contexts may have emerged in more complex settings. This hypothesis would address the fact that we predominately observed attenuation among negative interactions, as it would result in an apparent decrease in per-capita effect while actually representing an independent emergent positive effect.

A key finding here was that the relationship between individual isolate density and richness/total community density was informative for predicting the change in abundance of an isolate between community contexts (table 1). But why were changes in total density informative of changes in individual density? We suggest that this result arose because individual isolate density decreased as richness increased (figure 4b) due to the modest changes in total density (figure 4a). The associated attenuation of interactions in higher richness contexts was inherently observed as a decreased change in individual density but also a decreased change in total community density (figure 4c). This link between the two effects meant that the change in total density was an informative predictor of change in individual density (i.e., interaction effect) as well. Nonetheless, interaction effects themselves were useful predictors across contexts (table 2), suggesting that context-dependency generally does not redefine an interaction, but instead changes interactions to varying degrees. Indeed, we observed the explanatory power of interactions decayed as the divergence between community contexts increased (supplementary table 2). Such an outcome is in line with results from other studies, as it has been shown that predictions of coexistence based on pairwise cultures decay as complexity of the predicted community increases^23^.

We sought to advance our understanding of microbial interactions by observing how they vary across contexts and identifying patterns in that variation. Our observation of the general attenuation of interactions as richness increased is a straightforward and potentially useful result. And our finding that the relationships between individual density, richness and total density could help explain changes in pairwise interactions demonstrates both the usefulness of understanding community level properties and the value of considering interactions from the per-capita perspective. The observation that negative per-capita interactions nonetheless generally attenuated with richness suggests that context-dependency of interactions is a common feature in microbial communities. Further study of the specific processes that give rise to such context-dependence would be a fruitful endeavor that, combined with the observed population level processes, may improve our ability to predict the structure and function of microbial communities.

## Acknowledgements

We would like to thank members of the Bergelson laboratory for feedback, especially H. Maerkle for the inspiring discussions and constructive comments. This work was supported by an ERC Synergy grant, PATHOCOM (951444, J.B.) and the Hutchinson fund at The University of Chicago.

## Author Contributions

KD and JB conceived of the study and developed the study design. KD executed the experiments and performed data analysis, in consultation with JB. KD wrote the manuscript and designed the figures with editorial support and guidance from JB.

## Methods

### Resource availability

Lead contact: Keven D. Dooley

Materials availability: Bacterial genomes and experimental sequencing data generated in this study have been deposited to GenBank (Accession PRJNA953780).

### Data and code availability

All data reported in this paper will be shared by the lead contact upon request. All original code is available in this paper’s supplemental information.

Any additional information required to reanalyze the data reported in this paper is available from the lead contact upon request.

### Experimental Model Details

#### Bacterial strains

All bacterial strains were originally isolated from the leaves or roots of wild or field grown *Arabidopsis thaliana* in the midwestern states of the USA, specifically: IL, IN, MI (supplementary table 1). Strains were cultured at 28°C in a custom leaf-based culturing medium, “*Arabidopsis* leaf medium”.

#### *Arabidopsis* leaf medium (ALM)

*Arabidopsis thaliana* (KBS-Mac-74, accession 1741) plants were grown in the University of Chicago greenhouse in sterile potting soil at 50% humidity from January to March 2020. Seeds were densely planted in 15-cell planting trays, stratified for 3 days in the dark at 4°C, then moved to the greenhouse and thinned after germination to 4-5 plants per cell. Above ground plant material was harvested just before development of inflorescence stems. Plant material was coarsely shredded by hand before adding 100g to 400mL of 10mM MgSO_4_ and autoclaving for 55 minutes. After cooling to room temperature, the medium was filtered through 0.2μm polyethersulfone membrane filters to maintain sterility and remove plant material. The medium was stored in the dark at 4°C. Before being used for culturing, the medium was diluted 1:10 in 10mM MgSO_4_.

### Method Details

#### Experimental set up and culturing

Fresh bacterial stocks were prepared by first inoculating the isolates into 1mL of ALM shaking at 28°C and growing overnight. 100uL of these cultures were then used to inoculate 5mL of ALM shaking at 28°C. Once the cultures were visibly turbid, they were divided into 1mL aliquots with sterile DMSO added to a final concentration of 7% as a cryoprotectant. Stocks were stored at -80°C. Additionally at this time, stocks were diluted and plated to quantify density through colony counting.

To initiate an experiment, stocks were diluted to target densities determined by the initial community titer (∼1×10^6^ cells) and the number of initial members. For the preliminary synthetic communities, isolates were first combined into 7-member pools, subsequently combined into all 127 combinations of pools, and then distributed into three randomly selected wells containing 600μL of ALM in sterile 1mL deep-well plates. Similarly, for the synthetic communities used to measure interactions, isolates were first combined into desired initial community compositions and then randomly distributed in triplicate into 1mL deep-well plates. All such manipulations were performed under an open atmosphere with a Tecan Freedom Evo liquid handling robot. Deep-well plates were covered with sterilized, loosely fitting plastic lids to allow air exchange. Plates were cultured in the dark at 28°C on high-speed orbital shakers capable of establishing a vortex in the deep-well plates to ensure that the cultures were well-mixed. After 24 hours, 6μL of each culture was manually transferred by multi-channel pipette into new plates containing 594μL of fresh ALM. The new plates were immediately returned to the incubator and the day-old plates were stored at -80°C. The sample plates from the final time point (day 6) were amended with 15% glycerol prior to storage in the freezer to preserve the cultures for subsequent colony counting.

#### DNA Extraction

DNA was extracted from synthetic communities using an enzymatic digestion and bead-based purification. Cell lysis began by adding 250μL of lysozyme buffer (TE + 100mM NaCl + 1.4U/μL lysozyme) to 300μL of thawed sample and incubating at room temperature for 30 minutes. Next, 200μL of proteinase K buffer (TE + 100mM NaCl + 2% SDS + 1mg/mL proteinase K) was added. This solution was incubated at 55°C for 4 hours and mixed by inversion every 30 minutes. After extraction, the samples were cooled to room temperature before adding 220μL of 5M NaCl to precipitate the SDS. The samples were then centrifuged at 3000 RCF for 5 minutes to pellet the SDS. A Tecan Freedom Evo liquid handler was used to remove 600μL of supernatant. The liquid handler was then used to isolate and purify the DNA using SPRI beads prepared as previously described^45^. Briefly, samples were incubated with 200μL of SPRI beads for 5 minutes before separation on a magnetic plate, followed by two washes of freshly prepared 70% ethanol. Samples were then resuspended in 50μL ultrapure H2O, incubated for 5 minutes, separated on a magnetic plate, and supernatant was transferred to a clean PCR plate. Purified DNA was quantified using a Picogreen assay (ThermoFisher) and diluted to 0.5ng/μL with the aid of a liquid handler.

#### Sequencing library preparation

Libraries were prepared using Illumina Nextera XT kits and following a custom, scaled down protocol and custom indices (supplementary table 3). This protocol differed from the published protocol in two ways: 1) the tagmentation reaction was scaled down such that 1μL of purified DNA, diluted to 0.5ng/μL, was added to a solution of 1uL buffer + 0.5μL tagmentase, and 2) a KAPA HiFi PCR kit (Roche) was used to perform the amplification in place of the reagents included in the Nextera XT kit. PCR mastermix (per reaction) was composed of: 3μL 5X buffer, 0.45μL 10mM dNTPs, 1.5μL i5/i7 index adapters, respectively, 0.3μL polymerase, and 5.75μL ultrapure H2O. The PCR protocol was performed as follows: 3 minutes at 72 °; 13 cycles of 95 °C for 10 seconds, 55 °C for 30 seconds, 72 °C for 30 seconds; 5 minutes at 72 °C; hold at 10 °C. Sequencing libraries were manually purified by adding 15μL of SPRI beads and following the previously described approach, eluting into 12μL of ultrapure H2O. Libraries were quantified by Picogreen assay, and a subset of libraries were run on an Agilent 4200 TapeStation system to confirm that the fragment size distributions were of acceptable quality. The libraries were then diluted to a normalized concentration with the aid of a liquid handler and pooled. The pooled libraries were concentrated on a vacuum concentrator prior to size selection for a 300-600bp range on a Blue Pippin (Sage Science). The distribution of size-selected fragments was measured by TapeStation. Size-selected pool libraries were quantified by Picogreen assay and qPCR (KAPA Library Quantification Kit).

#### Sequencing

We characterized the compositions of our synthetic communities with a shallow metagenomics approach. We chose this approach as opposed to 16S amplicon sequencing as some of our isolates had identical 16S sequences and preliminary work with mock synthetic communities demonstrated that amplicon sequencing yielded less accurate characterizations of community composition. Reference genomes and initial synthetic community samples were sequenced on a HiSeq 4000 platform while follow up synthetic community samples were sequenced on a NovaSeq 6000 platform (paired end 2×150bp for both platforms). Reads were quality filtered and adapter/phiX sequences were removed using BBDuk from the BBTools suite^46^ (v38.81), with the following read quality parameters: qtrim=r, trimq=25, maq=25, minlen=50. Reads were mapped to reference genomes using Seal (BBTools) twice, once with the “ambig” flag set to “toss” (where ambiguously mapped reads were left out) and once with the “ambig” flag set to “random” (where ambiguously mapped reads were randomly distributed to equally likely references). By comparing the results between these two strategies, we identified sets of reference genomes which resulted in high numbers of ambiguous reads due to genomic similarity. We corrected for such ambiguity by reallocating “tossed” reads according to the proportion of unambiguous reads mapped to each isolate in the set for a given sample. To avoid mischaracterizing the composition of our synthetic communities due to contamination or non-specific mapping, for a given sample, isolates with less than 1% of total mapped reads were ignored.

#### Reference genome assembly

Reference genomes for the isolates used in these experiments were assembled using Spades v3.13.0^47^ with the “careful” flag. Assembled genomes were then manually curated in the Anvi’o^48^ software platform (v6.2), specifically using the interactive interface to remove outlier contigs assembled from contaminating sequences. The Anvi’o functions “anvi-summarize” and “anvi-estimate-scg-taxonomy” were also used to estimate the completion and contamination of assembled genomes and assign taxonomy based on single-copy core genes, respectively. The isolate names, taxonomy, and assembly information are presented in supplementary table 1.

#### Estimating absolute abundance

Absolute density of each community culture was measured by counting colonies from serial dilutions of the cultures. Specifically, glycerol preserved final timepoint samples were plated on 1X tryptic soy agar (TSA) plates, in triplicate serial dilution (3e^-5^, 1e^-6^, and 3e^-6^ dilutions), and cultured at room temperature. Colony forming units (CFU) were counted by eye over the course of the next few days. Final estimates of absolute abundance were calculated as the mean CFU/μL.

### Quantification and Statistical Analysis

#### Calculation of interactions

Population level effects were calculated as the ratio of focal isolate density with and without the interactor and presented as (ratio - 1) for ease of interpretation. The per-capita effects were calculated as the change in focal isolate density between contexts with and without the interactor, divided by the density of the interactor from the context in which it was present. To remove spurious interactions that arise from the presence of low abundance isolates close to the 1% relative abundance threshold, we pruned interactions to only include those within one standard deviation from the mean for both the population level and per-capita effect measures.

#### Statistical analysis and data visualization

Details of statistical tests are reported in the results section and figures. Statistical analysis and figure generation was performed in R^49^ v4.0.2 with aid from the following packages: tidyverse^50^ v1.3.0, reshape2^51^ v1.4.4, and car^52^ v3.0-11. Linear regression was performed with the “lm” function in R. All scripts are provided in the supplementary materials.

## Supplemental Item Titles

supplementary figure 1: Time course of community dynamics for an 8-member synthetic community

supplementary figure 2: Example of context-dependent coexistence

supplementary figure 3: Comparison of day-6 and day-12 compositions for 13 different communities assembled for measuring interactions

supplementary figure 4: Individual isolate density generally decreased as richness increased

supplementary table 1: Bacterial strain details

supplementary table 2: Summary of linear regressions modelling the predictive power of interactions across any richness contexts

Supplementary Table 3: Custom Illumina library indices

