## Supplemental Figures and Table for "Richness and density jointly determine context dependence in bacterial interactions"

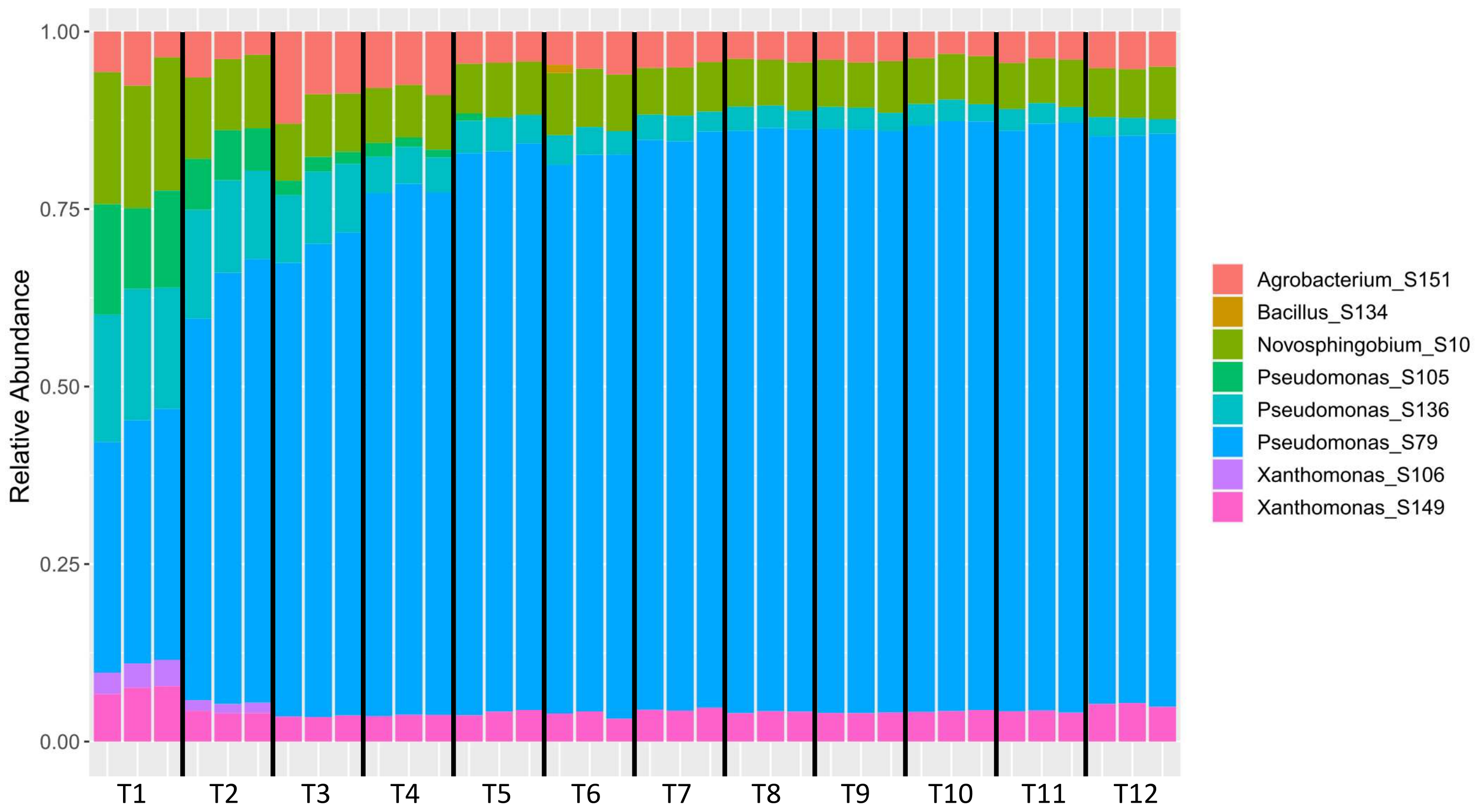


Supplementary Figure 1: A time course of community dynamics for an 8-member synthetic community. Each set of three stacked bars represents the composition of three replicates 24 hours after initial assembly or passaging for 12 days (T1 – T12). Dynamics play out quickly within the first four days and settle into a stable composition to until at least 12 days post initial assembly.

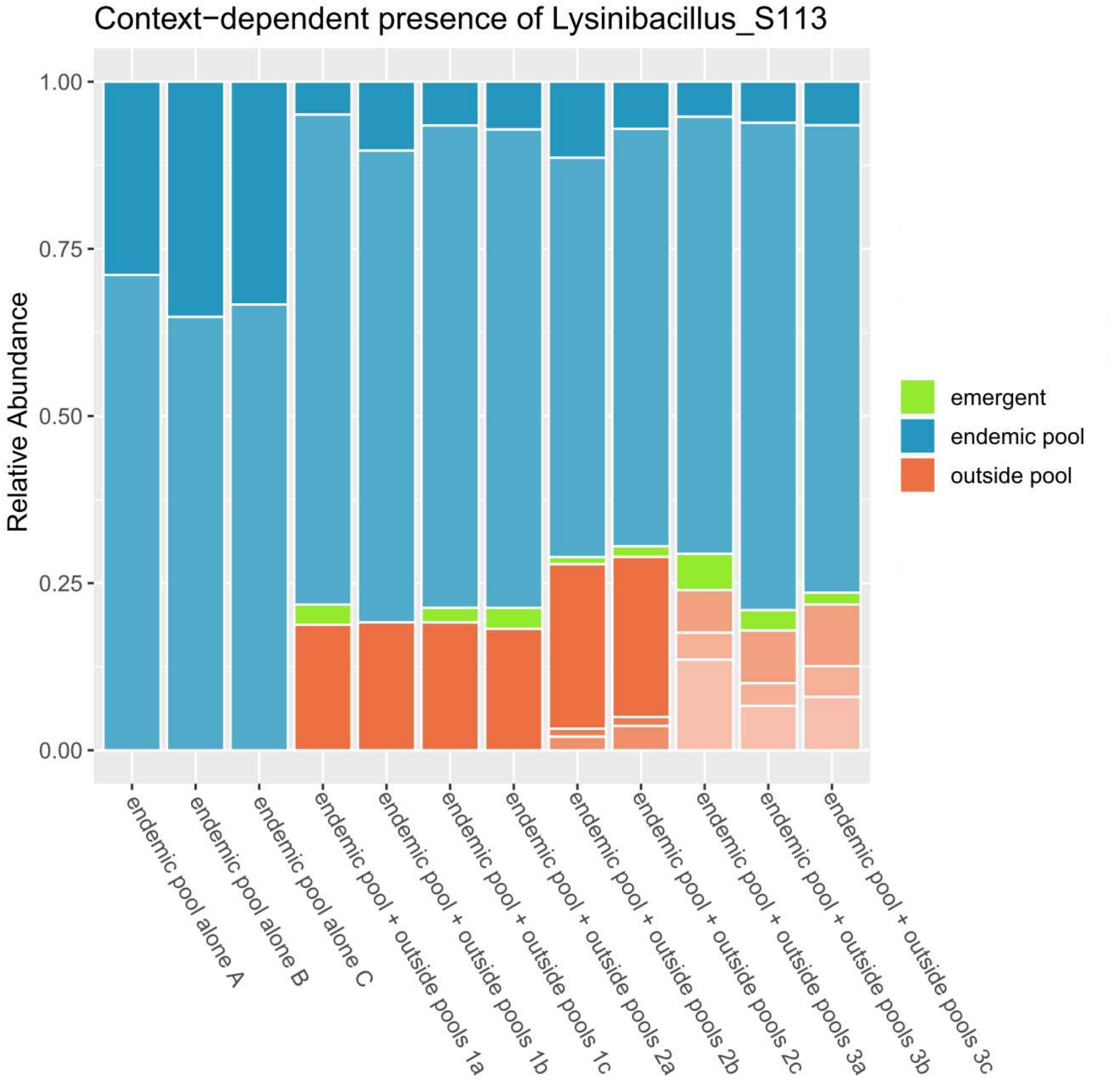


Supplementary Figure 2: An example of context-dependent coexistence of a Lysinibacillus isolate ("emergent" isolate) from the initial set of synthetic communities. That Lysinibacillus isolate was excluded by the other members of the 8-member pool to which it belonged ("endemic pool alone A-C"). However, in the three additional contexts shown here ("endemic pool + outside pools 1-3") which were assembled from that pool of 8 and at least one other pool, the Lysinibacillus isolate persisted to 6-days. The relative abundances of the non-emergent isolates are depicted by stacked bars in shades of blue (if from the "endemic pool" to which the Lysinibacillus isolate belonged) or shades of orange (if from pools other than the endemic pool).

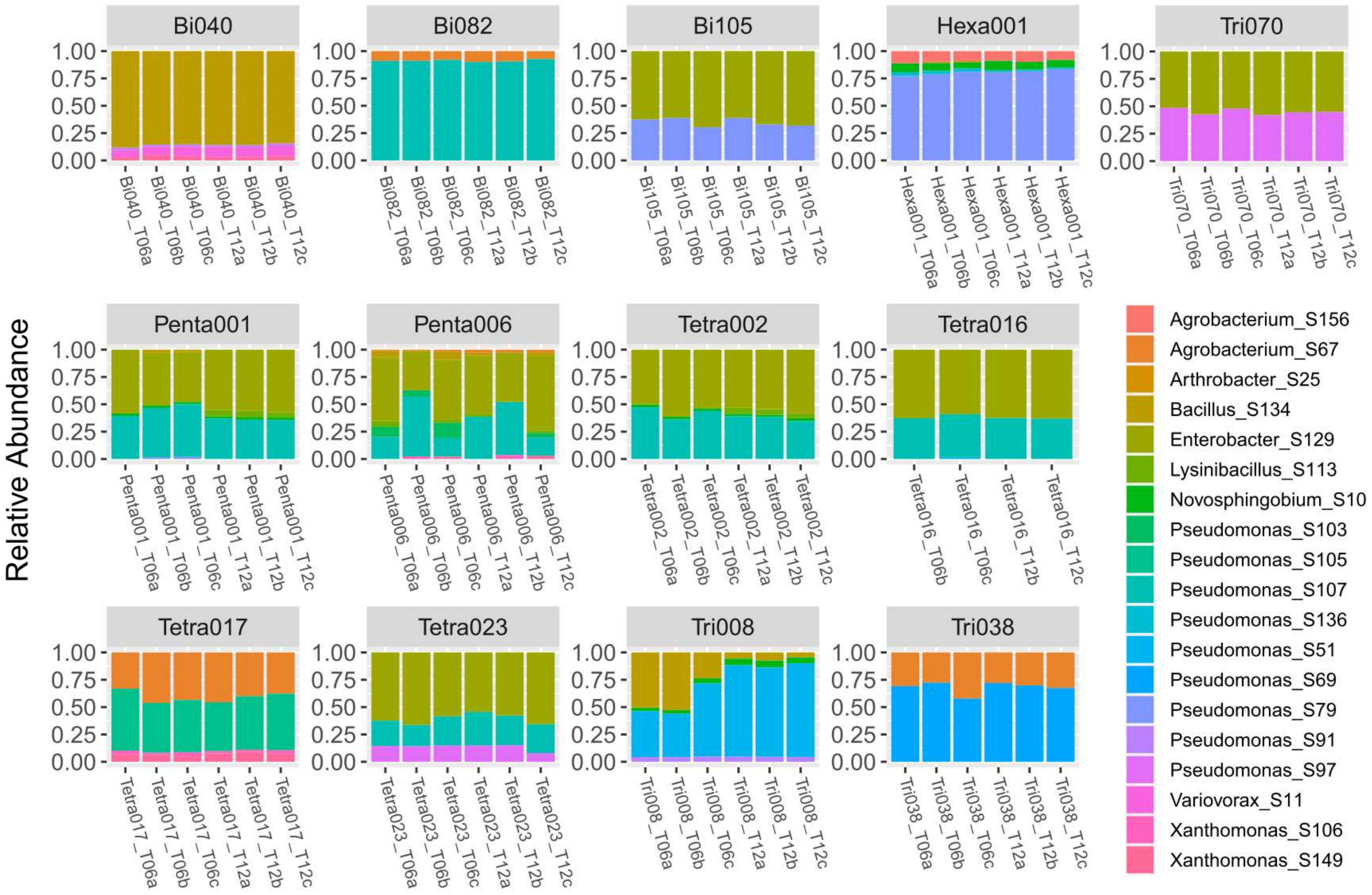


Supplementary Figure 3: A comparison of day-6 and day-12 compositions for 13 different communities assembled for measuring interactions. Timepoint is displayed in the x-axis text for each stacked bar. Stacked bars are colored by isolate identity. As seen in the full time-series (supplementary figure 1), community compositions after 6 days is generally representative of community compositions after 12 days. These communities were assembled from pools varying in initial richness (2-8 isolates), but all show generally consistent compositions between days 6 and 12, suggesting assemblages with higher initial richness do not require a longer period to reach a stable composition.

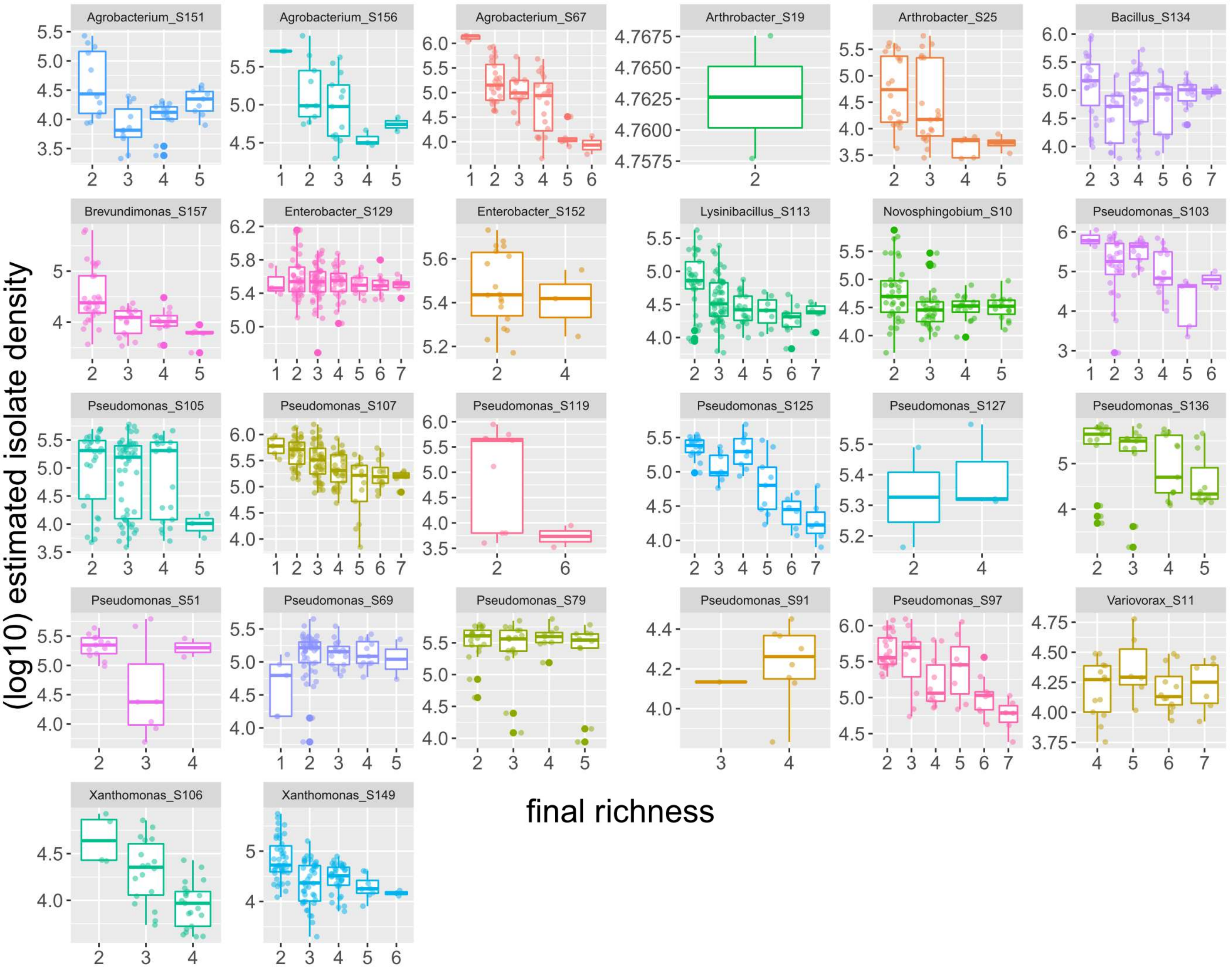


Supplementary Figure 4: Individual isolate density generally decreased as richness increased. Densities of each isolate at each observed richness is plotted on a log scale.

| model | df | adjusted $R^2$ | p-value |
|---|---|---|---|
| *all contexts* | | | |
| focal change ~ total change | 1 | 0.3521 | $< 2.2e^{-16}$ |
| focal change ~ interaction effect | 1 | 0.09991 | $3.428e^{-14}$ |
| focal change ~ total change + interaction effect | 2 | 0.4298 | $< 2.2e^{-16}$ |

Supplementary Table 2: Summary of linear regressions modelling the predictive power of interactions across any richness contexts (e.g., 1=>2 & 4=>5). “Focal change” indicates the change in density of the focal isolate in the predicted context. “Total change” indicates the change in total density between the community contexts of the interaction. “Interaction effect” indicates the change in density of the focal isolate in the interaction context which was not being predicted. We considered predictive power of interactions from the bottom-up, i.e., “interaction effects” came from the lower richness context (continuing the example above, 1=>2), while “total change” came from the predicted higher richness context (4=>5). A “+” in a model indicates the predictors were modeled as separate variables with no interaction.
